# Genome wide analysis of *Ga1-s* modifiers in maize

**DOI:** 10.1101/543264

**Authors:** Preston Hurst, Zhikai Liang, Christine Smith, Melinda Yerka, Brandi Sigmon, Oscar Rodriguez, James C Schnable

## Abstract

A one way reproductive barrier exists between most popcorn varieties and dent corn varieties grown in the United States. This barrier is predominantly controlled by the *ga1* locus. Using data from a diverse population of popcorn accessions pollinated by a dent corn tester, we found that the non-reciprocal pollination barrier conferred by *ga1* is more complex than previously described. Individual accessions ranged from 0% to 100% compatible with dent corn pollen. Using conventional genotyping-by-sequencing data from 371 popcorn accessions carrying *Ga1-s*, seven significant modifiers of dent pollen compatibility were identified on five chromosomes. One locus may either be a nonfunctional *ga1* allele present within popcorn, or second necessary gene for the reproductive barrier in genetic linkage with *ga1*, while the other modifiers are clearly genetically unlinked. The existence of *ga1* modifiers segregating in a popcorn genetic background may indicate selective pressure to allow gene flow between populations, which should be incorporated into future models of the impact of genetic incompatibility loci on gene flow in natural and agricultural plant populations.

## Introduction

Sexual incompatibility is a common phenomenon in the plant kingdom. Crossing barriers have been observed in several species, including tobacco^1, 2^, tomato^3–5^, and rice^6, 7^ among others. Lateral barriers, such that members of one population are unable to produce offspring with members of another population, can reinforce speciation^8^ and genes that create these barriers can play a critical role in cases of sympatric speciation^9, 10^. Due to the preferential and selective nature of the barrier, a male and female action are both required to enable recognition and discrimination of self and non-self pollen^11^. This type of incompatibility is seen in *Solanum*^12, 13^, as well as in some *Brassica* species^14^. It is possible that the action of both the male and female may be conferred through the actions of a protein encoded by a single gene, or the male and female functions of a crossing barrier may be provided by separate genes in tight genetic linkage, as has been shown for *tcb1*^15^.

In maize, the term gametophyte factor is employed to describe a locus where pollen grains carrying a particular allele exhibit increased fertilization success relative to pollen grains of the competing allele^16^. At least 10 such loci have been described for maize^16^. Of these *Ga1* was the first to be identified, with the first report of segregation distortion linked to the as yet unnamed *su1* locus on maize chromosome 4 published by Carl Correns in 1902^17^, the determination that *Ga1* was an independent genetic locus and not a second phenotype of the *su1* by R. A. Emerson in 1925^18^, and given the name *Ga1* by Mangelsdorf and Jones one year later^19^. *Ga1* creates a substantial unidirectional crossing barrier between plants carrying different alleles at the locus. Seed set when *Ga1-s* silks are pollinated with exclusively *ga1* pollen is consistently reported to be <5%^20, 21^, and in a competitive environment containing both *Ga1-s* and *ga1* pollen, estimates of the proportion of kernels successfully pollinated by *ga1* pollen grains range from 0.6-4%^19, 22, 23^. The strength of the *Ga1* crossing barrier is notable relative to other crossing barriers characterized in maize. The reference allele of *Ga2* allows 10-25% fertilization of wild-type pollen in competitive pollination assays, while a more recently isolated allele of *Ga2* from teosinte provides somewhat stronger incompatibility with wild-type pollen^24^. *Tcb1* provides near complete exclusion of *tcb1* pollen in a teosinte background, but exhibit attenuated effects when introgressed into maize backgrounds, suggesting the function of *Tcb1* depends on one or more modifier loci^25^. The early identification of *Ga1*, combined with the relatively extreme incompatibility phenotype of *Ga1* relative to other gametophyte factor loci described in maize^16^ have made the locus a particular focus of genetic investigation over the last century.

Three functionally distinct alleles have been described at the *Ga1* locus. The predominant allele in US maize germplasm is *ga1*. *Ga1-s* is found in many popcorn lines and in some populations of teosinte, the wild progenitor of maize, creates the unidirectional crossing barrier, as *Ga1-s* silks are receptive to pollen carrying the *Ga1-s* allele, but are largely non-receptive to pollen grains carrying the *ga1* allele^17, 26^. A third allele *Ga1-m*, conveys the male, but not female function of the *ga1* locus. Thus, pollen carrying *Ga1-m* can successfully pollinate *Ga1-s* silks, but *Ga1-m* silks can be successfully pollinated by *ga1, Ga1-m*, and *Ga1-s* pollen^27^. Data suggests the male function of *Ga1-m* may be incomplete, as *Ga1-s* pollen is still more successful than *Ga1-m* pollen when pollinating *Ga1-s* silks^28^. More recent studies suggest that *Ga1-m* may be the predominant allele in maize tropical landraces^29^.

Mapping *Ga1* proved challenging. Early maps based on phenotypic markers placed the locus on chr4 between *de1* and *su1* but at a substantial distance, perhaps 43 centimorgans (cM) from *de1* and 33 cM from *su1*^22^. Zhang and colleagues employed molecular markers to constrain the location of *Ga1* to a 1.5 cM interval on chromosome 4^20^. Employing additional data from the B73 x Hp301 NAM subpopulation and a backcross population carrying Ga1-s, Bloom and Holland constrained the location of *ga1* to a location containing 13 predicted genes in the B73 reference genome^21^. Lauter et al.(2017) provided evidence PME3, a pectin methylesterase, is responsible for the female action of the locus^30^. Zhang et al.(2018) identified a PME gene as responsible for the male action of *Ga1-s*^31^. These studies suggest that both the male and female actions of unilateral dent sterility are related to the deesterification of pectin. Recently another pectin methylesterase *Pertunda* has also been identifed as responsible for the female action of the *Tcb1* crossing barrier^32^, although pollen carrying the *Tcb1-s* and *ga1* alleles is compatible with *Ga1-s* silks, and vice versa.

Mapping receptitivity to *ga1* pollen in a popcorn by dent corn recombinant inbred line population (B73 x Hp301) identified only a single locus corresponding to *Ga1*^21^. Fine mapping estimated that locus to be slightly larger than 1MB^33^. In contrast to the result from the B73 x Hp301 population, apparent modifiers of *ga1* function have been reported in the popcorn variety Supergold^34^, from which many of the inbreds from one the three major heterotic groups in commercial hybrid popcorn production are derived^35^. Here we employed data from *ga1* pollination tests of a diverse panel of maize popcorns, including representatives of the Supergold, Amber Pearl, and South American popcorn heterotic groups to assess the degree of variation present for *Ga1* and whether specific modifiers segregating in popcorn could be identified.

## Methods

### Scoring Dent Sterility

A diverse panel of 311 popcorn lines planted in an isolation plot at the University of Nebraska Experimental Station at Mead, NE during the 2017 mid-western United States growing season were assessed for quantitative compatibility with *ga1* pollen. Each genotype was represented by 20 plants planted in a 17.5 foot long row, with a 30 inch alley between each range and each row. The perimeter of the field was planted with two rows of a purple *ga1* line in a dent corn background. Within the field, every fifth row was planted with the same purple dent *ga1* line. Lines of the diversity panel were detasseled prior to anthesis and female flowering dates were recorded for each accession to confirm exposure of the silks to freshly shed pollen from the purple dent *ga1* line. Each line was scored for incompatibility with *ga1* pollen at maturity. Ears with significant numbers of yellow or white kernels, indicating pollen contamination from a source either than the intended *ga1* pollen donor were disregarded. (Figure S1)

### Genotyping

327 commercial inbred popcorn lines from the ConAgra Foods popcorn breeding program and 44 public popcorn lines from USDA were genotyped using conventional genotyping-by-sequencing (GBS)^36^. DNA samples were submitted to the Cornell Institute for Genomic Diversity for GBS library construction and sequencing. Samples were sequenced in four distinct barcoded libraries. A total of 827M raw Illumina reads were generated. Reads were aligned to the B73 reference genome (RefGen V3) using bwa aln^37^. 238,772 SNPs were called using the TASSEL-GBS pipeline v3.0^38^ with the MAF (minor allele frequency) filtering threshold set to 0.05 and mnF (minimum inbreeding coefficient defined as 1-Ho/He) set to 0.8. Missing SNP calls were imputed using Beagle version 4.1^39^. After imputation, SNPs with a minor allele frequency below 0.01 were removed from the dataset. The final dataset consisted of 238,772 SNPs scored in 311 maize popcorn lines for which both genotypic and phenotypic data was available.

### Genome Wide Association Analysis and Interpretation

Association analyses were performed in parallel using the mixed linear model (MLM) implemented in GEMMA with a significance threshold of 0.01^40^ and the FarmCPU algorithm (a multi-locus mixed model (MLMM)) implemented in MVP (https://github.com/XiaoleiLiuBio/rMVP) at a Bonferroni corrected significance threshold of 0.01^41^.

The first 5 principal components of population structure, as determined by TASSEL^42^, were including in both analysis to control for population structure effects^43^. Kinship was calculated using MVP and the same kinship matrix was employed in both analyses. (Figure S2) Structural and functional gene annotations for version 3 of the B73 reference genome were obtained from Phytozome v12.1. All genes within 10kb of the SNP were considered potential candidate genes. All linkage disequilibrium calculations were performed in TASSEL, version 5^42^.

## Results

Significant variation in the degree of *ga1* fertilized seed set was observed among the accessions of this popcorn population (Figure S1). Half of accessions showed >5% seed set with purple kernels indicating successful pollination by *ga1* pollen. Five lines showed near complete compatibility with *ga1* pollen. This observation is notably different from reports which assessed *Ga1-s* introgressed into dent corn backgrounds where Ga1-s consistently produces a high degree (>95%) of sterility when fertilized with *ga1* pollen^20, 21^.

A genome-wide association study was conducted to test whether specific loci outside the *Ga1-S* locus were associated with variation in the effectiveness of the *Ga1-s* allele’s female function. Using genotype calls for 238,772 segregating SNPs genotyped using conventional genotyping by sequencing, seven statistically significant trait associated SNPs were identified on five chromosomes using the FarmCPU algorithm: 2, 3, 4, 5 and 10 (Figures 1, S2). Three of these trait associated SNPs (Chr5 bp:99960879, Chr4 bp:7020954, and Chr2 bp:72824565) were also supported by MLM based GWAS (Figures 1, S3). The trait associated SNP identified on chromosome 4 is located at 7,020,954 bp on the B73_RefGenV3 reference genome, corresponding to approximately 7.8MB in B73_RefGenV4. This position is in close proximity to two genes identified at the *Ga1* locus^30, 31^.

The precise placement of the *Ga1* locus in the B73 reference genome remains somewhat ambiguous. Multiple groups who have attempted to map the *Ga1* region have only been able to narrow it down to a large interval^20, 21, 31, 33^, likely as a result of structural differences between the popcorn derived *Ga1-s* and dent corn derived *ga1* haplotypes at this locus. Consistent with this explanation, three large gaps containing no SNP markers of size 429kb, 826kb, and 454kb were observed within the estimated position of the *Ga1-s* locus, in comparison to 1 SNP every 12kb in the whole dataset (Figure 3). Here we employ the interval reported by Zhang et al which placed the causal gene between 8.52MB and 10.23MB on chromosome 4 in B73 RefGenV4. After translation, the equivalent is approximately 7.66Mb to 9.35Mb on chromosome 4 in B73 RefGenV3^31^. The range of the gaps found in the current study span from 7.623Mb to 9.358Mb.

**Figure 1.**
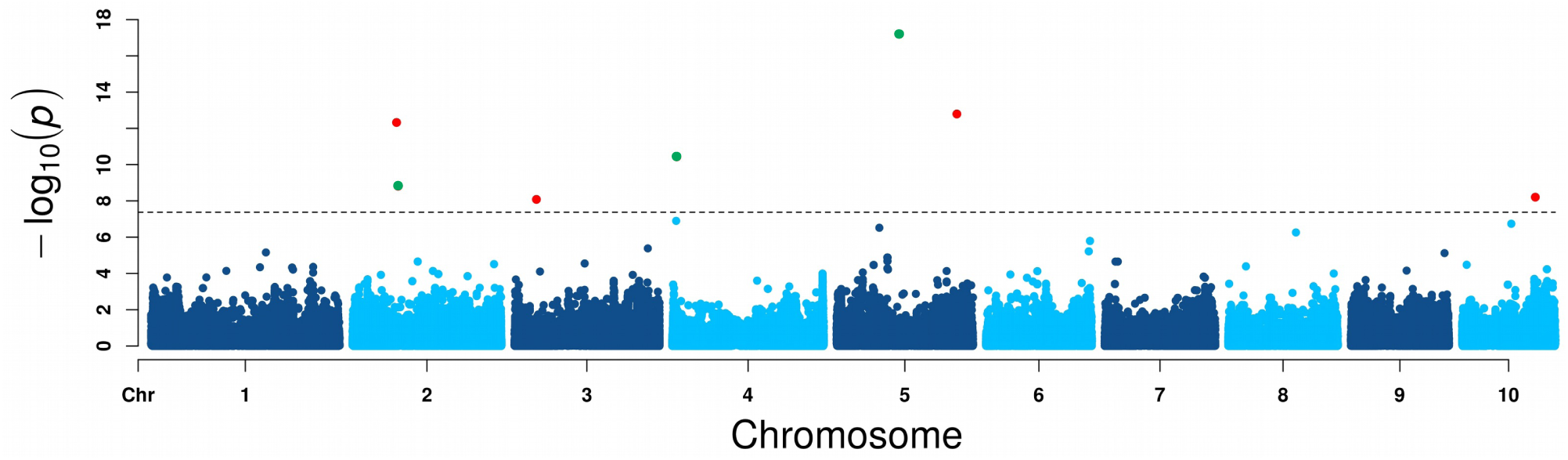
Associations between 238,772 genetic markers and degree of receptivity to *ga1* pollen using data from 311 popcorn lines. Each point indicates a single SNP, and position on the y-axis indicates the statistical significance of the association between that marker and sterility/fertility as determined using FarmCPU. Dashed line indicates a threshold of p=4.18e-8 (bonferroni corrected threshold based on the number of SNPs tested and an original threshold of p = 0.01). For SNPs above the bonferroni corrected threshold, larger green points were also statistically significant in a parallel MLM GWAS analysis using the same genetic and phenotypic data.

The distribution of phenotypic scores for individuals carrying the minor allele at the chr4 locus are also consistent with the minor allele of this trait associated SNP marking a *ga1* allele, either from introgression of dent germplasm during popcorn breeding and improvement, or already present at low frequencies in popcorn germplasm. However, based on patterns of linkage disequilibrium in the region from 6.7Mb to 10.5Mb on chromosome 4 (Figure 3) it appears that the trait associated SNP Chr4 bp:7020954 is in not in linkage with the region from 8.53Mb to 10.23Mb in the B73 reference genome determined to correspond to the popcorn region carrying the pectin methylesterases responsible for the *ga1* reproductive barrier^31^. In addition, Chr4 bp:7020954 exhibits significant LD with adjacent markers in the genome, indicating the marker is likely not misplaced, whether as a result of a misassembly in the B73 reference genome or structural variation between popcorn and dent corn (Figure 3).

The minor alleles of the trait associated SNPs identified on chromosomes 2 and 5 were also associated with an increase in successful fertilization by *ga1* pollen (i.e. decreased effectiveness of the reproductive barrier conveyed by *Ga1-s*). The minor alleles of trait associated SNPs on chromosomes 3 and 10 are associated with a decrease in successful fertilization by *ga1* pollen (i.e. increased function of *Ga1-s* as a reproductive barrier (Figure 2). Three trait associated SNPs (Chr5 bp:192080965, Chr4 bp:7020954, Chr10 bp:117590335) are within larger intervals identified by composite interval mapping^44^. Composite interval mapping also identified a signal on chromosome 2 that, while not sufficiently strong to be statistically significant on its own, overlaps with and lends additional confidence to the two trait associated SNPs (Chr2 bp:70339953 and Chr2 bp:72824565) identified on that chromosome^44^. With two exceptions, Chr2 bp:72824565 and Chr5 bp:192080965, each trait associated SNP was within 10KB of one or more annotated maize gene models (Table **??**). However, caution should be used in interpreting these gene lists as significant presence absence variation is present in maize^45, 46^ and genes involved in reproductive barriers in popcorn may be absent from the B73 maize reference genome as was, indeed, the case for both *Ga1-s* and *Tcb1-s*^30–32^.

**Figure 2.**
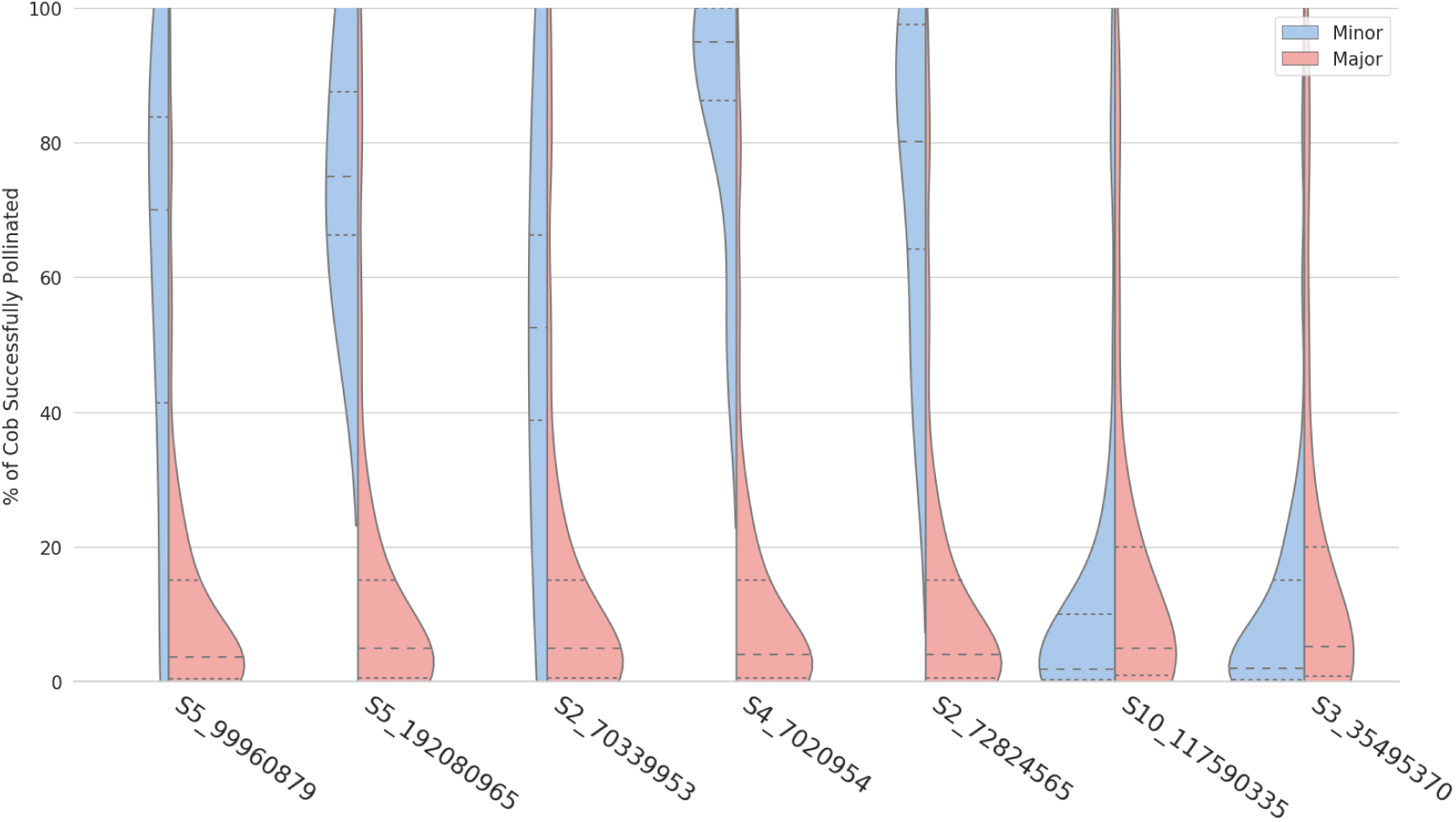
Distribution of % receptivity to *ga1* pollen for individuals carrying either the minor or major alleles of each of the seven significantly trait associated SNPs identified in the FarmCPU GWAS analysis. Trait associated SNPs are ordered from most significant p-value (left) to least significant p-value (right). Lines with longer dashes indicate the median trait value for each subpopulation. Lines with shorter dashes indicate the 25th and 75th percentiles of the trait distribution for each subpopulation.

## Discussion

The results presented above suggest the genetic basis of non-reciprocal cross sterility conferred at the *Ga1* locus is more complex than previously thought. Anecdotally, popcorn breeders have long noticed a complex mode of inheritance^34^, and at least one previous study identified mapping intervals which appeared to act as modifiers of *Ga1-s*^44^. The existence of these modifiers pose challenges to efforts to employ the *Ga1-s* allele in commercial breeding programs.

One trait associated SNP, Chr4 bp:7020954, is in close proximity to the *ga1* region, and may indeed represent a *ga1* allele, either from introgression of dent germplasm during popcorn breeding and improvement, or already present at low frequencies in popcorn germplasm. However linkage disequilibrium analysis was not consistent with this interpretation (Figure 3). One alternative explanation is that Chr4 bp:7020954 marks a second gene in tight linkage with the PME genes already identified as essential to *Ga1-s* function which modifies the function of this crossing barrier. GRMZM2G157241, located 8kb from Chr4 bp:7020954, is homologous to characterized calcium binding proteins. It has been proposed that transfer of calcium ions formed from the action of pectin methylesterase enzymes plays a role in the Ga1 locus^31^. As pectin is deesterified by PME, free Ca2+ becomes crosslinked with pectin and the cell wall of the pollen tube becomes stiff^47^. However further research will be needed to determine whether Chr4 bp:7020954 indeed marks a tightly linked modifier of *Ga1-s* or, despite the present linkage disequilibrium evidence, marks a knockout of at least the female function of *Ga1-s* equivalent to either *Ga1-m* or *ga1*.

**Figure 3.**
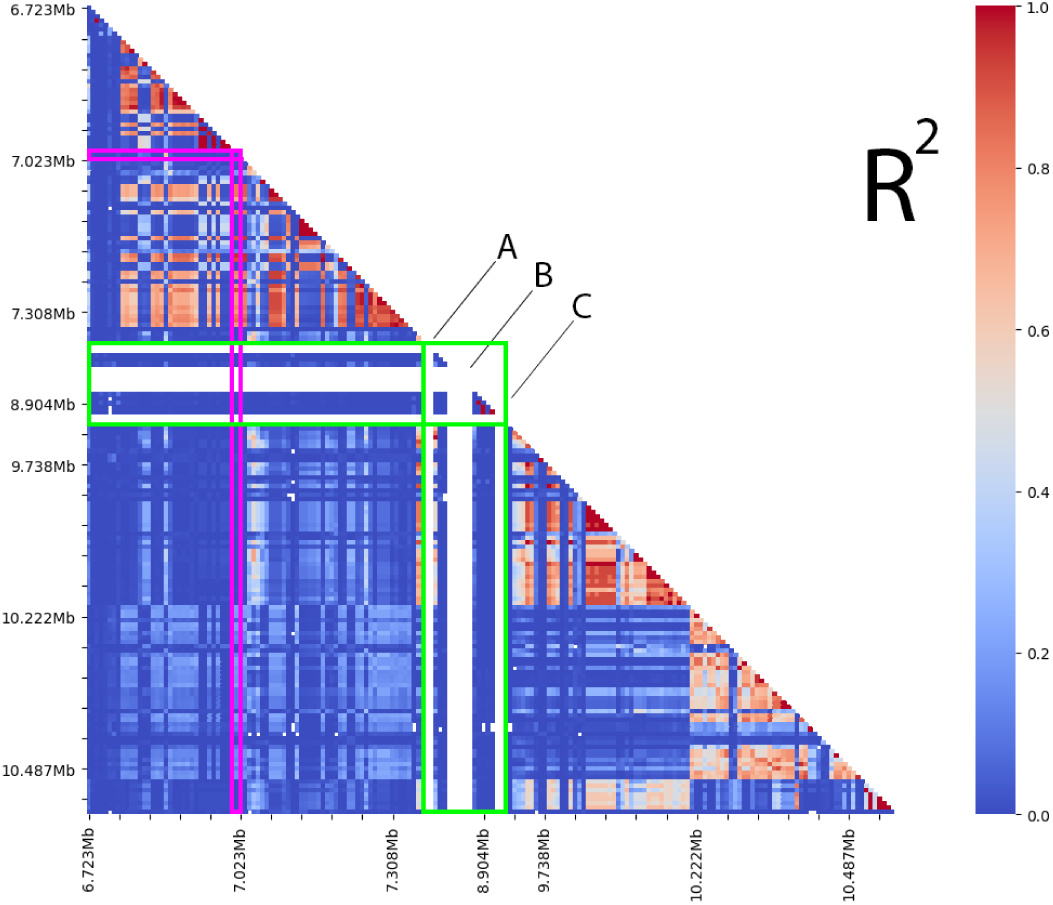
Linkage disequilibrium among SNP markers between 6.7Mb to 10.5Mb on chromosome 4. Each row/column represents a single marker. The purple outline marks the trait associated SNP S_7020954. The green outline marks the interval for *Ga1-s* reported by Zhang et al. 2018, translated from B73 RefGenV4 to B73 RefGenV3. Three large gaps without any SNP markers identified in this population, are indicated by the white gaps – not drawn to scale – labeled A, B, and C. **A**, is a gap without any markers from 7.623Mb to 8.052Mb; **B**, is a gap without any markers from 8.078Mb to 8.904Mb; **C**, is a gap without any markers from 8.904Mb to 9.358Mb.

**Table 1.**
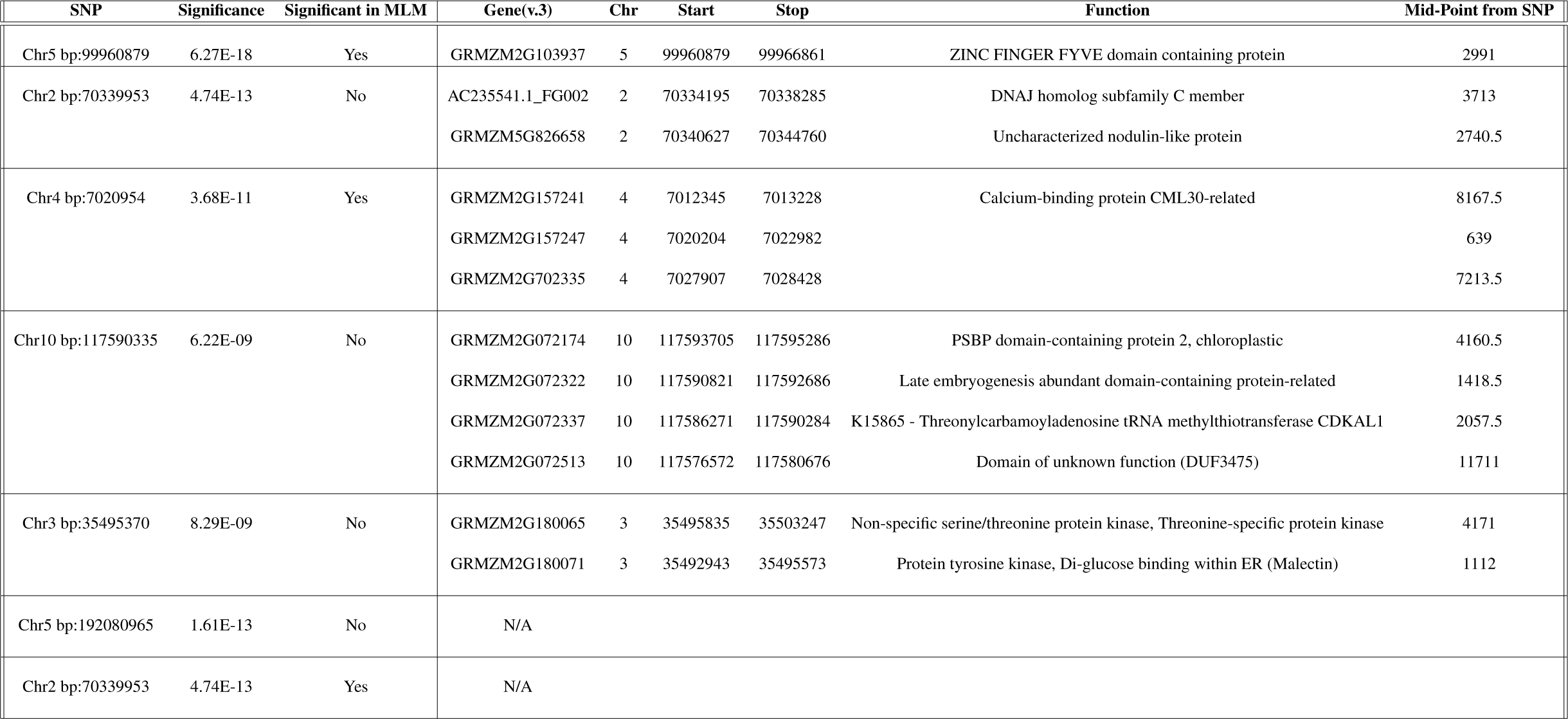
Table 1: All genes within 10KB of significant SNPs.

The existence of modifiers for *Ga1-s* function also has implications for specialty maize breeding and production efforts. One recent proposal has been to introgress the *Ga1-s* allele into maize refuge lines and grow in a mixed bag refuge system with Bt maize^48^. As the Bt *ga1* pollen would be unable to pollinate the ears of the non-Bt *Ga1-s* refuge plants, in theory this approach would increase the amount of non-Bt grain available to insect pests, decreasing the selective pressure for Bt resistance in pest species. *Ga1-s* modifers present a challenge and an opportunity, as *ga1* pollen is likely to dramatically outnumber *Ga1-s* pollen on the silks of refuge plants within Bt fields, and this any increase or decrease in *Ga1-s* efficiency is likely to produce substantial shifts in the effectiveness of this refuge strategy. Some organic producers also utilize the *Ga1-s* allele as an additional approach to avoiding any detectable GM traits from pollen drift which might threaten organic certification^49^. Here again, knowledge of loci that increase or decrease the effectiveness of the crossing barrier could have significant economic impact.

If allelic differences impact the degree to which populations inter-mate, it will surely have an impact on allele frequencies in other areas of the genome and impact the amount of gene flow from one population to another. If speciation genes are subject to modification by other loci, one would expect that all loci involved would have an impact on the history and future of the organism. It is possible that selective sweeps exist around these loci, that may explain some of the phenotypic variation between wild and domesticated genotypes, as well as different classes of maize such as dent, flint and popcorn.

## Author contributions

BS, JCS, MY and OR conceived the study; CS, OR, and ZL generated the data, PH and ZL analyzed the results, JCS and PH wrote the paper

## Acknowledgements

Supported by research funding from ConAgra Brands to JCS and OR. The authors thank Aaron Lorenz and Amritpal Singh for their assistance in popcorn genotyping.

---

**Figure S1.**
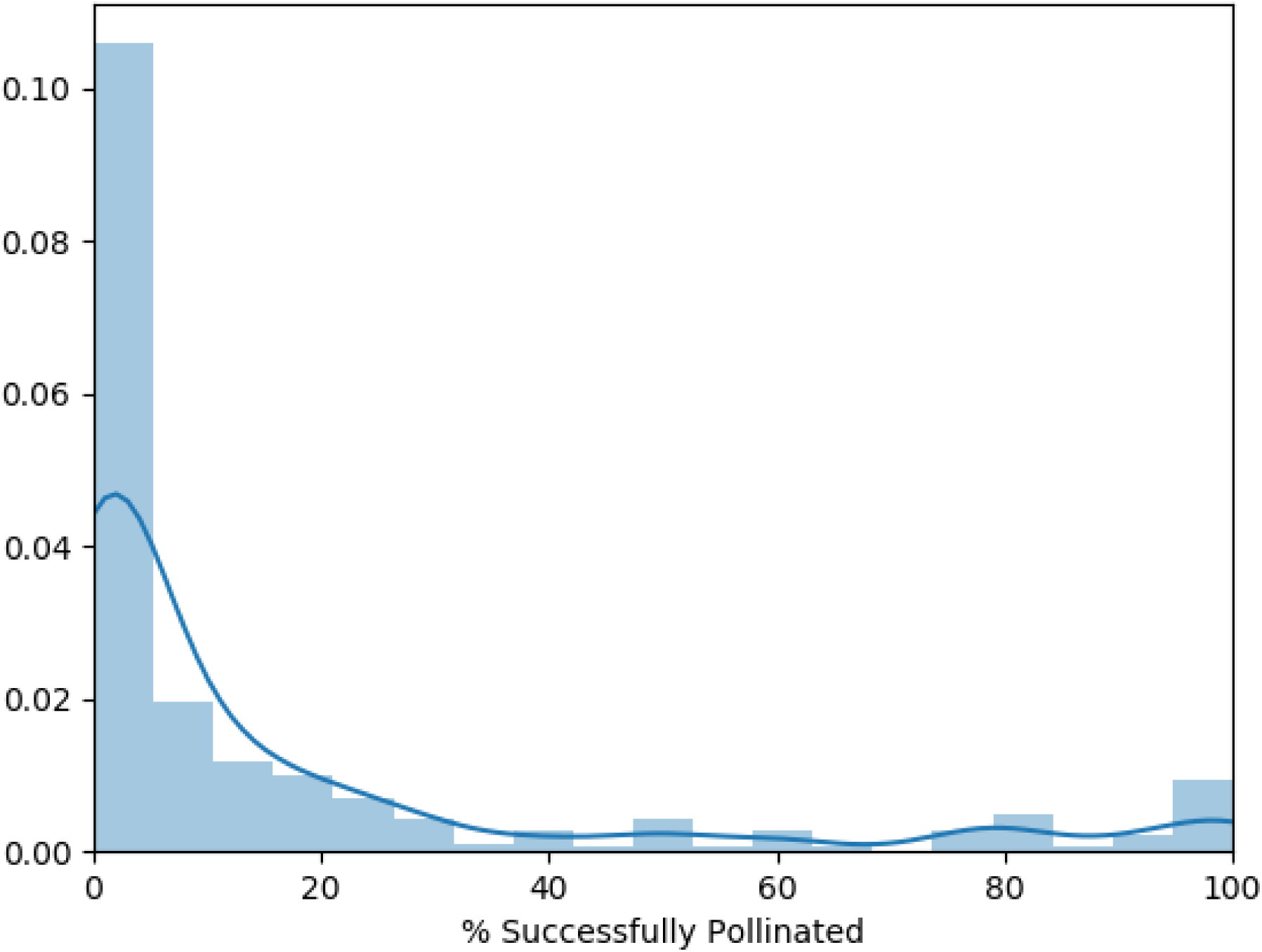
Distribution of percent compatibility with *ga1* pollen among the 311 popcorn lines employed in this study. Let’s remove the tick marks from the bottom of the histogram -James

**Figure S2.**
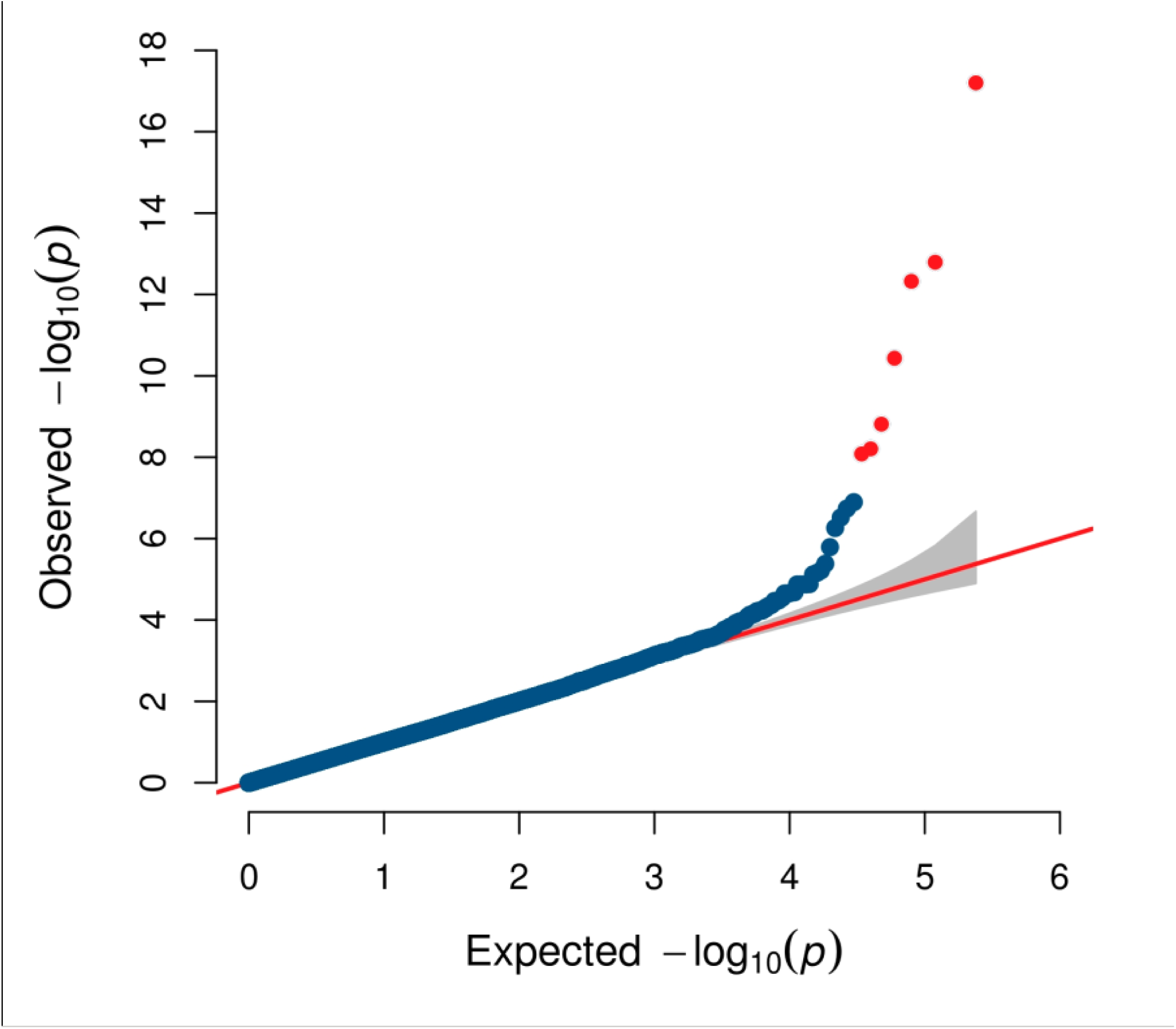
Overrepresentation of significant p-values in the FarmCPU analysis relative to the distribution expected from random data. Points in red indicate p-values which considered to be statistically significant in this analysis (based on a bonferroni corrected p-value threshold of 0.01)

**Figure S3.**
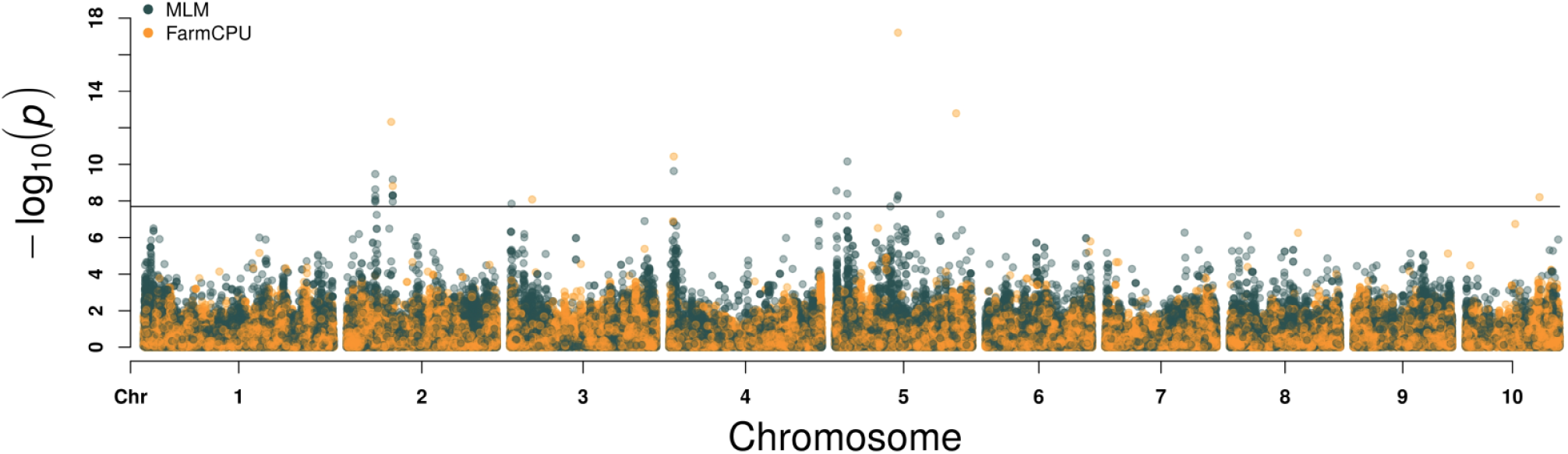
Combined results from both MLM and FarmCPU GWAS mapping. Each SNP is indicated by two separate points one indicating the p-value for the association between that SNP and variation in *ga1* pollen compatibility in the MLM GWAS analysis (green) and a second indicating the p-value for the association between that SNP and variation in *ga1* pollen compatibility in the FarmCPU GWAS analysis (green)

